# Beyond Natural Antibodies: Scaffold-Based Generation of Novel Anti-3CL^pro^ Nanobody Nb01

**DOI:** 10.1101/2025.05.06.652338

**Authors:** Zhijie Zhan, Jun Xiong, Yini Jiang, Yinghua Li, Yuanzhe Cai, Chong Luo, Feijuang Huang, Jieren Liu

## Abstract

In the post-pandemic era, continuous mutations and persistent infections of Severe Acute Respiratory Syndrome Coronavirus 2 (SARS-CoV-2) pose a significant threat to global health, making the normalization of preventive measures imperative. Due to its high conservation, the 3CLpro is relatively stable across all these variants without significant changes, suggesting that drugs targeting this enzyme could be effective against all variant viruses. And compared to traditional antibodies, nanobodies show significant advantages against SARS-CoV-2 and its variants. Traditional antibodies often lose their inhibitory activity due to viral mutation. Nanobodies are characterised by their small size, high stability, high affinity for antigen binding, and high water solubility, which enable them to capture viruses quickly and effectively address the continuous mutations and persistent infections of SARS-CoV-2. In this research, we devised a strategy to produce nanobodies by utilizing a fragment-generating large language model, resulting in the identification of a nanobody named Nb01. Nb01 exhibits potent and broad-spectrum neutralizing activity against numerous SARS-CoV-2 variants.

The nanobody Nb01, directed against the SARS-CoV-2 3CL^pro^, demonstrated efficacy against the majority of prevalent SARS-CoV-2 variants. Notably, it has a higher affinity than the current best performing nanobodies (S43, bn03, R14, and 3-2A2-4) for most of the variants, including the Alpha (27.5%), Gamma (29.7%), Omicron BA.2 (32.2%), BA.4/5 (81.2%), BF.7 (64.2%) and XBB (5.5%) variants. For other variants, Nb01 displayed affinities that were on par with these benchmark nanobodies. In summary, the exceptional specificity, low toxicity, robust stability, and extensive spectrum of Nb01 indicate its potential to be developed as a nanobody therapeutic for the management of SARS-CoV-2 infections and its diverse variants.

## I. Introduction

In the post-pandemic era, while there is a gradual resumption of societal activities, the normalization of preventive measures remains imperative [22, 16]. The persistent evolution of the SARS-CoV-2 virus has resulted in the continual appearance of novel variants, most notably the Omicron variants and its sub-variants BA.4 and BA.5 [27]. These variants demonstrate increased transmissibility and infectivity, and immune evasion [48], thereby posing a substantial threat to global public health [34]. The substantial changes in their antigenic properties have facilitated these VOCs (Alpha, Beta, Gamma, Delta, and Omicron) to escape from serum neutralization of convalescent and vaccinated individuals, leading to an increase in new infections among the unvaccinated but also break-through infections among the infected and vaccinated individuals [26].Concurrently, persistent SARS-CoV-2 infection [19], commonly referred to as Long COVID or Post-Acute Sequelae of SARS-CoV-2 infection (PASC), has emerged as a novel public health challenge.The persistent SARS-CoV-2 infection often goes unrecognised, and therefore might affect a substantial number of people, particularly immunocompromised individuals. Furthermore, in the setting of suboptimal immune responses,such persistent infections could lead to the emergence of new variants that evade immunity on the individual and population levels [30]. Despite significant progress in vaccination, uncertainty about the virus continues to make global health a serious challenge.

SARS-CoV-2 undergoes continuous mutation and evolution, giving rise to different variants. However, a specific proteinase within the virus, 3-chymotrypsin-like protease (3CL^pro^), remains relatively stable across all these variants without significant changes [32, 7]. Major protease (3CL^pro^) is a highly conserved enzyme of SARS-CoV-2. The enzyme plays pivotal role in the virus replication cycle [37, 29].Enzyme stability is vital for the biological function of the virus, and also means that it could be an ideal target for drug development, as drugs targeting this enzyme may be effective against all variants [3, 17, 1, 13]. And compared to traditional antibodies, nanobodies show significant advantages against SARS-CoV-2 and its variants [6].Traditional antibodies, although highly specific, tend to lose their inhibitory activity due to Virus mutation [14, 15].Nanobodies are characterised by their small size, high stability, high affinity for antigen binding and high water solubility, which enable them to capture viruses quickly and efficiently stop pathogens from spreading [35].The stable structure and suitable mechanism of action of nanobodies make them a superior therapeutic option to traditional antibodies, offering a broader targeting capability and effectively addressing the continuous mutation and persistent infection of SARS-CoV-2 [6, 5].

Nanobodies have showcased remarkable potential in countering SARS-CoV-2 and its variants, yet their development has encountered significant hurdles, notably the challenge of generating effective nanobody structures in their entirety (Fig. 1) [24]. These sequences, which is directly generated by fine-tuning of ProtGPT2 through a nanobody library, fail to properly recognize the loop structure and beta-barrel structure of nanobodies, since the unique structural properties and functions of nanobodies [20]. Characterized by beta-barrel structures and loop structures, these unique features set nanobodies apart from conventional proteins [33]. Beta-barrel structures, composed of multiple beta-strands, confer a robust structural foundation to nanobodies. The loop structures, particularly the cyclic variable regions known as complementarity-determining regions (CDRs), are pivotal in antigen binding and exhibit high sequence variability, allowing them to accommodate a diverse array of antigenic epitopes. These structural hallmarks endow nanobodies with the ability to bind a wide spectrum of antigens with exceptional stability. However, the intricate and high-dimensional nature of this structural data poses a challenge for neural networks, which struggle to capture the profound relationships and patterns within it [11, 45]. The complexity of modeling how amino acid sequences fold into the distinctive beta-barrel and loop configurations is beyond the current capabilities of these networks. This limitation hampers the accurate prediction and generation of the overall nanobody structure, thereby constraining the advancement of nanobody-based therapeutics.

**Fig. 1.**
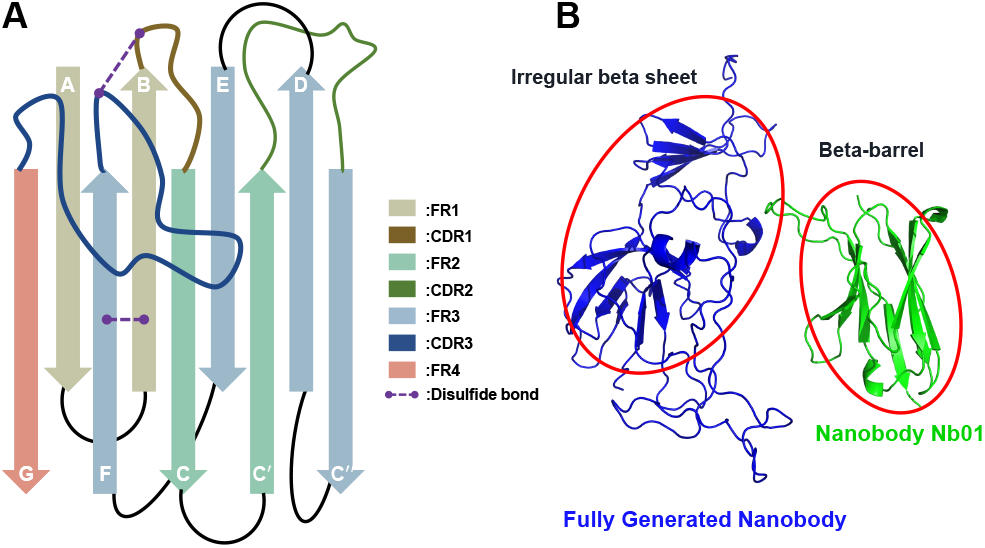
Analysis of nanobody structures. (A) The general structure of nanobodies [33]; (B) The protein structure directly generated by ProtGPT2 vs. the correct protein structure of nanobody (Nb01).

### Observation

We analyzed the sequence structure of the nanobodies. The results (see Fig. 2) indicated that the amino acid diversity score was significantly higher in the complementary determining region (CDR), with diversity indices ranging from 0.2 to 0.6 [9], compared to the framework region (FR), which had a diversity index below 0.2. Moreover, in a biological sense, CDRs are the variable regions of antibodies; they contain the amino acid residues most closely associated with antigen binding and determine the specificity and affinity of the antibody [43]. Framework Region (FR) is the region adjacent to the CDR in the variable region of an antibody that contains the framework structure and support system of the antibody [28]. Although the direct contribution of FRs to antigen binding is relatively small, they play an important role in the stability of antibodies, the formation of antibody structure, and the regulation of antibody function. Based on these findings, we identified the FR regions as conserved regions of the anti-SARS-CoV-2 nanobodies, which could be utilized for segmental training of ProtGPT2 to generate CDR sequences for the development of novel nanobodies.

**Fig. 2.**
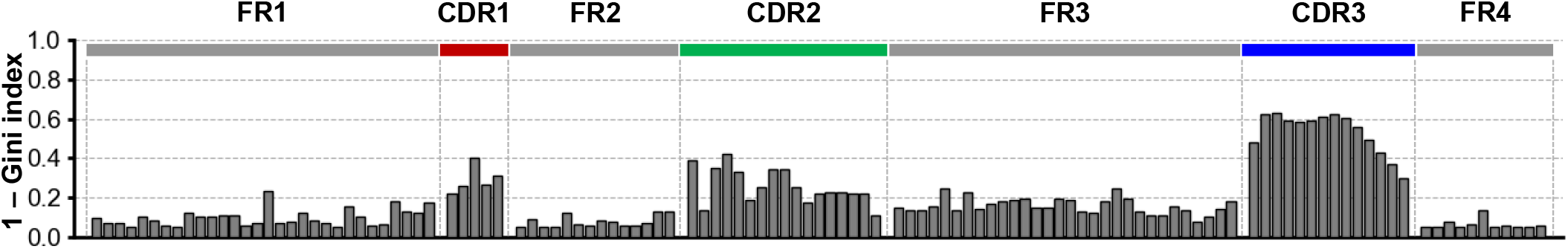
Diversity score (1-Gini Index) for amino acid sites in nanobody libraries. A diversity score (*≤*0.2) is generally indicated that this site is conserved [8].

### Solution

Considering the high diversity of CDRs and the conserved nature of FRs, we designed a framework for the generation of nanobodies against SARS-CoV-2 3CL^pro^. That is, we apply ProtGPT2 to generate CDR sequences and then graft them with FR regions to obtain novel nanobodies against SARS-CoV-2. Our Scaffold-based generated method consists of the following steps: (i) building a nanobody library; (ii) training to generate CDR sequences; (iii) selection of the widely used humanized framework h-NbBcII10_FGLA_ as a scaffold; (iv) clustering and compressing CDR sequences, and randomly selecting compressed sequences to be grafted to the scaffolds; (v) predicting the structure of the nanobodies using Alphafold2, and (vi) evaluating the characterization of the nanobodies. In the experimental part, the nanobody Nb01, directed against the SARS-CoV-2 3CL^pro^, demonstrated efficacy against the majority of prevalent SARS-CoV-2 variants. Notably, it exhibited the highest affinity for the Alpha, Gamma, Lambda, Omicron BA.2, BA.4/5, BF.7, and XBB variants, surpassing the current top-performing nanobodies, including S43, bn03, R14, and 3-2A2-4.

### A summary of our contributions

- **Deep learning-based nanobody development framework**: we designed a Scaffold-based deep learning generation method and successfully developed an effective anti-3CL^pro^ nanobody Nb01 we designed a Scaffold-based deep learning generation method and successfully developed an effective anti-3CL^pro^ nanobody Nb01.
- **The nanobody Nb01 has superior physicochemical properties**: The GRAVY (Grand Average of Hydropathicity), Toxins, Theoretical pI, Instability Index and Aliphatic Index of Nb01 were calculated to be −0.17, 0.41, respectively, 7.94, 41.52 and 72.22, respectively, and were characterized by good solubility, low toxicity, high stability, and a significant content of aliphatic acids. The physicochemical properties of Nb01 are superior compared to the existing well-known nanobody S43.
- **The nanobody Nb01 is able to effectively target a wide range of variants**: The nanobody Nb01, directed against the SARS-CoV-2 3CL^pro^, demonstrated efficacy against the majority of prevalent SARS-CoV-2 variants. Notably, it has a higher affinity than the current best performing nanobodies (S43, bn03, R14, and 3-2A2-4) for most of the variants, including the Alpha (27.5%), Gamma (29.7%), Omicron BA.2 (32.2%), BA.4/5 (81.2%), BF.7 (64.2%) and XBB (5.5%) variants.

## II. Method

Fig. 3 depicts a schematic diagram of the main steps of this study, which includes: (i) constructing of an anti-SARS-CoV-2 nanobody library (see Section II-B) (ii) generating sequences in CDR area using ProtGPT2 fine-tuned (see Section II-C); (iii) selecting of the widely used humanized framework h-NbBcII10_FGLA_ as a scaffold (see Section II-D); (iv) clustering of sequence in CDR to omit redundant sequences and randomly selecting non-redundant CDR sequences grafting with the scaffold to build new nanobodies (see Section II-E); (v) predicting the structure of the nanobodies using Alphafold2 (see Section II-F) and (vi) evaluating the properties of these generating nanobodies (see Section II-G).

**Fig. 3.**
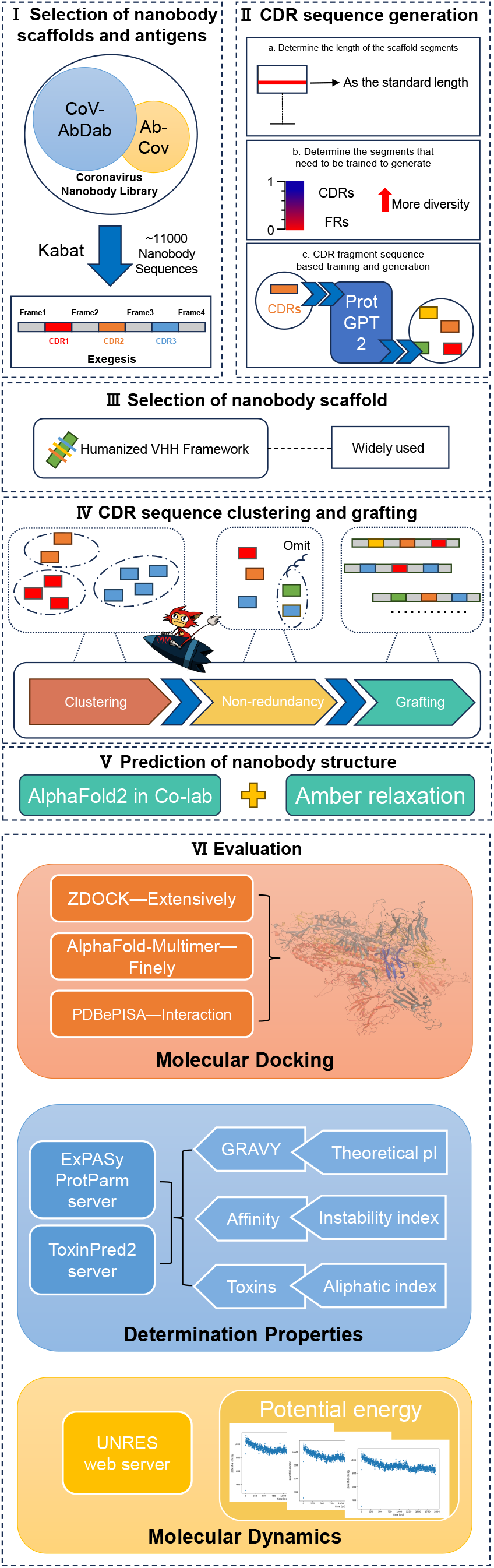
A schematic diagram for the main steps conducted in this study.

### A. Task Description

Our task is to generate novel efficient anti-pan-SARS-Cov-2 nanobodies targeting 3CL^pro^. We also evaluate a series of properties of the nanobodies and validated the effectiveness of the generated sequences.

### B. Library construction and analysis of coronavirus nanobodies

Nanobody sequences were downloaded from CoV-AbDab [39] and Ab-Cov [38], and 11,000 coronavirus nanobodies were collected (see Table I). Nanobodies were separated into CDRs and framework regions (FRs) using Kabat annotation of AbRSA [25]. The region between FR1 and FR2 is CDR1, the region between FR2 and FR3 is CDR2, and the region between FR3 and FR4 is CDR3.

**TABLE I.** Data set characteristics.

| Data set | Antibody | Nanobody | Nanobody average length |
| --- | --- | --- | --- |
| CoV-AbDab | 12536 | 11981 | 122.82 |
| Ab-CoV | 1963 | 1766 | 122.25 |

### C. CDR sequence generation

First, the standard length of each segment in the nanobody library was determined. Second, the diversity index was calculated and combined with the biological significance of each segment to select the segments that needed to be trained for generation. In the end, we used ProtGPT2 to generate segments.

#### 1) Determine the length of the scaffold segments

The lengths of CDRs and frames (FRs) in the database were analyzed, and the appropriate lengths were selected as the canonical number of amino acids for CDRs and frames (FRs), and CDRs or frames (FRs) smaller than the canonical number of amino acids were populated with empty position holders to the same number. We chose the median of the individual fragments as the respective fragment lengths, so that the standard lengths of FR1, CDR1, FR2, CDR2, FR3, CDR3, and FR4 were 30, 5, 14, 17, 30, 12, and 11, respectively.

#### 2) Determine the segments that needed to be trained to generate

The level of diversity (diversity score) at each amino acid site was measured using the 1-Gini index [9]. The diversity scores take values between 0 and 1, where 0 indicates perfect homogeneity (100% abundance of amino acid) and 1 indicates the highest diversity (all amino acids have the same abundance). The diversity of each site is shown in Fig. 2.

#### 3) Model inference generation

To explore new CDRs after we delineated the antibody sequence data, we fine-tuned the ProtGPT2 model [12] by using the data of CDR1, CDR2, and CDR3 from CoV-AbDab and Ab-Cov data sets, respectively. The parameters of ProtGPT2 for fine-tuning are shown as follows. In addition to this, we also attempted to fine-tune ProtGPT2 using the sequences of the intact nanobodies.

### D. Selection of nanobody scaffold

In order to avoid the influence of the framework region on antibody activity, we chose the widely used humanized nanobody framework (h-NbBcII10_FGLA_) [46] as the basis for the nanobody design activity. Table II lists the corresponding FR scaffold fragments of h-NbBcII10_FGLA_.

**TABLE 2.** Sequence of scaffold frames (h-NbBcII10_FGLA_)

| Annotation Name | Amino Acid Sequences |
| --- | --- |
| FR1 | QVQLVESGGGLVQPGGSLRLSCAAS |
| FR2 | WFRQAPGQGLEAVA |
| FR3 | RFTISRDNSKNTLYLQMNSLRAEDTAVYYCAA |
| FR4 | WGQGTLVTVSS |

### E. CDR sequence clustering and grafting

CDR sequences were checked for redundancy and CDR sequences with high similarity were omitted. Nanobodies were obtained using non-used sequences grafted to standard frameworks.

#### 1) CDR sequence clustering

The CDR sequences were checked for redundancy and CDR sequences with high similarity were omitted. The CDR sequences in the clustering centers are all non-redundant CDRs and do not consume additional computational cost. We used MMseqs2 to cluster CDR sequences (with default parameters) [31, 42] and used the probability of a non-redundant CDR being selected as the ratio of the cluster size of the sequence in which it is located to the total set of sequences as the probability of selecting that non-redundant CDR.

#### 2) CDR sequence grafting

The new nanobodies were obtained by substitution of the selected CDR sequences with the corresponding standard frame sequences and then splicing with the scaffold frame sequences. In the nanobody library, non-redundant sequences are randomly selected and grafted with the standard framework to obtain the control CDR set (1000 nanobodies). In the generated CDRs, non-redundant sequences are randomly selected and grafted to the standard framework to obtain the generated CDR set (1000 nanobodies).

### F. Prediction of nanobody structure

The AlphaFold2 structures in this study were mainly generated using the AlphaFold2 implementation in the ColabFold notebooks [44] running on Google Colaboratory, using the default settings with Amber relaxation.

### G. Evaluation

#### 1) Evaluation pan-SARS-CoV-2 antigen

To analyze the broad-spectrum of anti-SARS-CoV-2, we select the main COVID-19 strains, such as SARS-CoV-2 Alpha (PDB ID: 7NEH), Beta (PDB ID: 7S5Q), Delta (PDB ID: 8R80), Lambda (PDB ID: 7U28), Gamma (PDB ID: 7NXC), Omicron BA.2 (PDB ID: 7ZF8), Omicron BA.4/5 (PDB ID: 7ZXU), Omicron BF.7 (PDB ID: 8XZD), Omicron BQ.1.1 (PDB ID: 8IF2), and Omicron XBB (PDB ID: 8XE9), in our experiments. Their structural files are downloaded from the RCSB PDB [4].

#### 2) Compared nanobody sequences

##### Fully generated nanobody sequences

Sequences directly generated by fine-tuning of ProtGPT2 through a nanobody library.

##### Scaffold-based generated sequences

The CDR1, CDR2, and CDR3 sequences generated by fine-tuning ProtGPT2, and then grafted to the h-NbBcII10FGLA framework.

##### Famous anti-pan-SARS-CoV-2 nanobody sequences (baseline)

We selected currently recognized nanobodies against each variant of SARS-CoV-2 as a baseline. (i) R14 and S43 [27]: It has potent anti-pan-SARS-CoV-2 virus activity and also displays a variety of pan-sarbecovirus activities with a nanobody structure derived from PDB ID: 7WD1 (R14) and 7WD2 (S43); (ii) 3-2A2-4 [26]: It is one of the most widespread and potent nanobodies described to date, with potency against Omicron subvariants, SARS-CoV-1 and key representative coronaviruses, with a nanobody structure derived from PDB ID: 7X2L; (iii) bn03 [22]: It is a highly potent bispecific single domain antibody with broad neutralizing ability against SARS-CoV-2 and its variants and can be administered by inhalation with a nanobody structure derived from PDB ID: 8I4E.

#### 3) Evaluation Metric

##### Distinguishing between nanobodies and non-nanobodies

To determine whether a protein sequence is a nanobody, we analyze the similarity score between the generated sequence and the consensus sequences of the nanobody based on AbRSA [25]. The similarity (%) metric [25] is a value ranging from 0 to 100 that measures how similar a sequence is to the consensus nanobody sequence: when the similarity is *>* 70, it is considered to be a nanobody; when the similarity is between 60 and 70, it is considered to be indeterminate; and when the similarity is *<* 60, it is considered not to be a nanobody.

##### Molecular docking

In order to understand the interaction of nanobodies with antigens, we performed two types of molecular docking: (i) *Wide-scale docking*: Extensive docking of antibody-antigen molecules using ZDOCK [36], which facilitates the calculation of the affinity distribution of the nanobodies and the screening of high-affinity antibodies, with default parameters used for the ZDOCK docking method; (ii) *Fine docking*: For a more precise understanding of antibody-antigen interactions, we used AlphaFold-Multimer [21] for further molecular docking, and the final file was optimized with the Amber force field. Further interaction analysis was performed using PDBePISA [23]; molecular visualization of the docking results was performed using PyMol (PyMOL Molecular Graphics System, Schrödinger, LLC). In summary, these data predicted differences in the binding epitopes of the nanobodies to the selected antigens as well as the docking results.

##### Prediction of physicochemical properties of nanobodies

To further analyze the distribution of physicochemical properties of the antibodies. The physicochemical properties of the nanobodies such as theoretical pI, the aliphatic index, the instability index and the grand average hydrophilicity (GRAVY) were evaluated using ExPASy ProtParam server [47]. Among them, theoretical pI was used to understand the behavior of proteins in different pH environments, aliphatic index to predict the thermal stability of proteins, instability index to assess the tendency of proteins to degrade in vivo or in vitro environments and GRAVY to analyze the hydrophilicity of proteins. Toxicity of nanobodies was assessed using Toxin-Pred2 server (default parameters) [41].

##### Molecular dynamics (MD) simulation

To further analyze the stability of nanobodies. Molecular dynamics (MD) simulations of nanobodies were performed using the UNRES online server (default parameters) [10] to analyze the dynamic stability of nanobodies. We performed MD runs to predict the potential energy at 300 K.

## III. Experimental results

In Section III, we (i) first distinguish whether the fully generated sequences and scaffold-based nanobodies are nanobodies; perform large-scale docking of nanobody libraries and scaffold-based nanobodies to nanobody antigens, respectively; and perform predictive analyses of physicochemical properties of nanobody libraries and scaffold-based nanobodies; (ii) fine docking of excellent scaffold-based nanobodies and baselines to nanobody antigens, respectively; analyzing the dynamic properties of nanobody-antigen complexes by molecular dynamics simulations.

### A. Scaffold-based generated sequences are nearly equal to the real nanobodies

#### 1) Scaffold-based generated sequences are much closer to real nanobodies and have a better originality than fully generated sequences

Fig. 4A shows the statistical results of the similarity between the different generated nanobody sequences and the consensus nanobody sequences. The similarity score of scaffold-based generated sequences is greater than **92.3** (*>* 80, nanobody), but the similarity value of the fully generated sequences is less than **38.3** (*<* 60, non-nanobody). This observation clearly indicates that because of the insufficient nanobody data sets (around 13 thousand anti– SARS-CoV-2 nanodoby), directly training and generating the nanobody sequences is not a correct way, but based on the stable scaffold, scaffold-based generated method yields the better results.

**Fig. 4.**
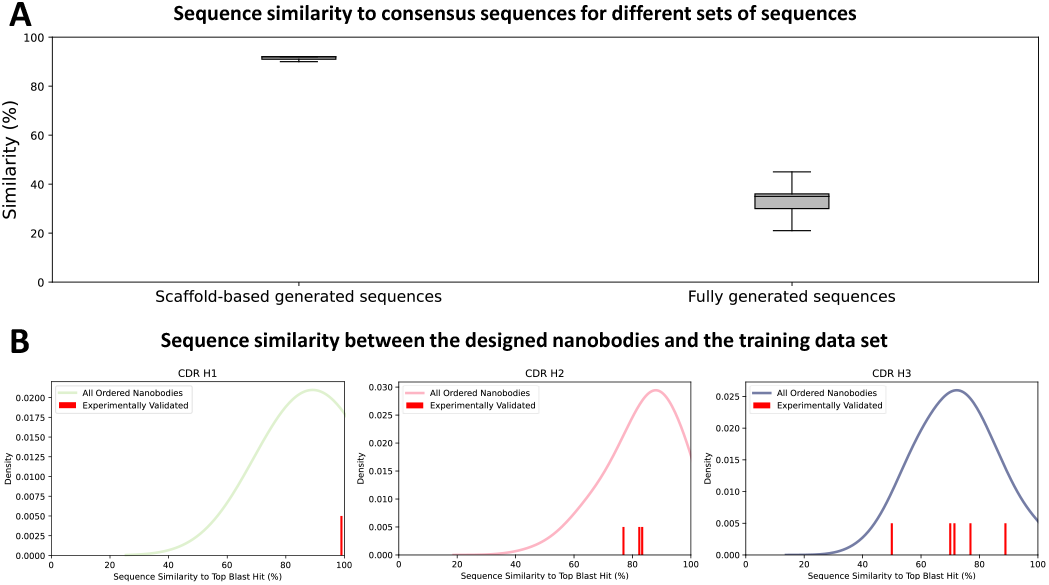
Sequence evaluation of designed nanobodies. (A) Similarity of the two sets of sequences to the consensus sequence of the nanobody (the evaluation principle of similarity score is shown in section II-G3); (B) Blastp [2] was used to find hits for CDR fragments based on the nanobody library.

##### Scaffold-based generation yields novel nanobody sequences

Fig. 4B shows the sequence similarity between scaffold-based generated sequences and existed anti-SARS-CoV-2 nanobody library (see Section II-B), and the peak of CDR1, CDR2, and CDR3 hits 90%, 85%, and 75%, respectively. Clearly, the diversity in CDR3 (see Fig 2) leads to generate more novel sequences in CDR3.

#### 2) The affinity of scaffold-based generated sequences is superior to that of nanobody library, and other properties are close to the real world nanobody library

##### The affinity of scaffold-based generated sequences for individual SARS-CoV-2 variants is generally better than that of current real-world effective nanobodies

We calculated the affinity activity and physicochemical properties of control CDR and generated CDR nanobodies. A comparison of the affinity distributions of control CDR and generated CDR nanobodies for each SARS-CoV-2 variant is shown in Fig. 5A. Higher ZDOCK Score indicates higher affinity. In terms of affinity, the affinity of scaffold-based generated sequences for each variant was overall better than that of nanobody data set because the median and box of scaffold-based generated sequences were both higher than that of nanobody library. In addition, we calculated the lower bound (LB) values of ZDOCK score for both sets of antibodies as a measure of the overall affinity of the nanobodies for SARS-CoV-2 variants. The formula for LB value is given below:

**Fig. 5.**
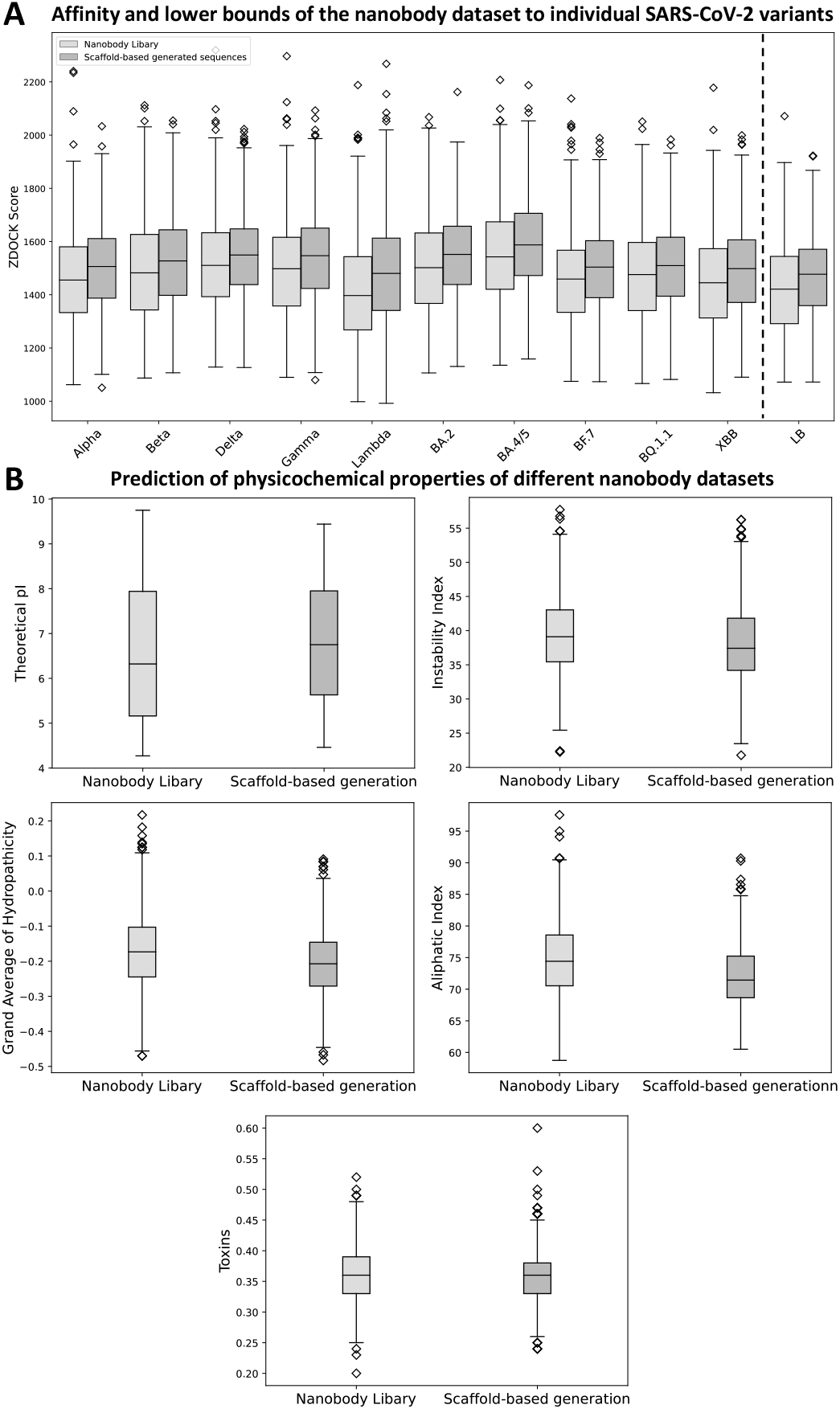
Property predictions for two sequence sets. (A) predictions of the distribution of affinity (ZDOCK Score) properties for individual SARS-CoV-2 variants; (B) predictions of the distribution of other physicochemical properties (Theoretical pI, Grand Average of Hydropathicity (GRAVY), Instability Index, Aliphatic Index, and Toxins).

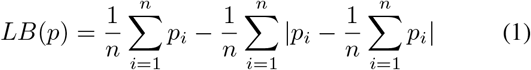

Where *p* is a set of nanobodies, *LB*(*p*) is the possible lower bound of *p, n* is the number of variants, and *p*_*i*_ is the affinity of *p* for the ith variant. This method does not depend on a specific distribution, is less sensitive to outliers, and is more suitable for small datasets (here *n* in our experiment is equal to 10) [18]. It can be noticed that in Fig. 5A, the LB value of ZDOCK score for scaffold-based generated sequences is overall 2.89% (41/1410) higher than that of nanobody library, indicating that scaffold-based generated sequences have more broad-spectrum SARS-CoV-2 affinity than existed nanobodies in library.

##### Scaffold-based generated sequences have the same physicochemical properties as nanobody libraries

Fig. 5B shows the predicted distribution of other physicochemical parameters (Theoretical pI, Grand Average of Hydropathicity (GRAVY), Instability Index, Aliphatic Index, and Toxins). The Theoretical pI, Grand Average of Hydropathicity (GRAVY), Instability Index, Aliphatic Index and toxicity of scaffold-based grerated sequences is equal to 6.8, *>* −0.3, *<* 45, *>* 65, *<* 0.6, respectively. This indicated that scaffold-based generated sequences have high hydrophobicity, high stability, high thermal stability and low toxicity. Meanwhile, in Table III, the JS divergence between nanobody libraries and scaffold-based generated sequences is less than 0.1, which shows that they have the similar distribution. We selected the five (Top5: Nb01-05) scaffold-based generated sequences with the highest lower bounds of affinity (LB) for the following experimental analysis. It should also be noted that the five nanobodies that were verified by Blastp to bind to the target are not similar to the current existed nanobodies (red line in Fig. 4B).

**TABLE 3.** JS divergence.

| Property | GRAVY | Toxins | Theoretical pI | Instability Index | Aliphatic Index |
| --- | --- | --- | --- | --- | --- |
| JS divergence | 0.0767 | 0.03734 | 0.0615 | 0.0611 | 0.0214 |

### B. Nb01 has excellent anti-pan-SARS-CoV-2 properties

#### 1) Nb01 has excellent physicochemical properties

Table IV shows the predicted physicochemical properties of Nb01 and the baseline. For aliphatic index, Nb01 is higher than all baselines, indicating that Nb01 has superior thermal stability; for instability index, the value of Nb01 is in the middle of the baselines; for GRAVY, the highest value of Nb01 indicates that Nb01 is more hydrophobic; for toxins, Nb01’s value is less than 0.6, which means the toxic is extremely low.

**TABLE IV.** Physicochemical properties of Nb01 and baseline.

| Name | Theoretical pI | Aliphatic Index (↑) | Instability Index (↓) | GRAVY (↑) | Toxins (↓) |
| --- | --- | --- | --- | --- | --- |
| Nb01 | 7.94 | 72.22 | 41.52 | -0.17 | 0.41 |
| 3-2A2-4 | 6.64 | 69.51 | 38.75 | -0.48 | 0.31 |
| bn03 | 4.69 | 69.08 | 47.49 | -0.21 | 0.41 |
| R14 | 6.36 | 58.54 | 45.04 | -0.41 | 0.31 |
| S43 | 5.06 | 53.06 | 41.50 | -0.60 | 0.39 |

#### 2) Nb01 has excellent binding affinity

*Nb01 is with the highest affinity for the three variants (Alpha, BA*.*4/5, and BF*.*7) and the second or third highest affinity for the other variants*. Table V shows the binding affinity of Top-5 and baseline to each SARS-CoV-2 variant. Meanwhile, to show the binding affinity for the same pocket of SARS-CoV-2, we do the competitive experiment between Nb01 and S43. Table VI shows that Nb01 is more competitive than S43 in eight in ten SARS-CoV-2 variants.

**TABLE V.** Affinity (kcal/mol) of the generated nanomers (lower bound Top5) and the baseline to the mutant, respectively.

| Name | Alpha | Beta | Delta | Gamma | Lambda | BA.2 | BA.4/5 | BF.7 | BQ.1.1 | XBB |
| --- | --- | --- | --- | --- | --- | --- | --- | --- | --- | --- |
| Nb01 | **-15.3** | -7.7 | -7.4 | **-14.4** | -8.8 | **-12.3** | **-12.5** | **-13.3** | -10.7 | **-11.5** |
| Nb02 | -15.1 | -9.3 | -7.3 | **-17.1** | -12.1 | -8.3 | -11.9 | -6.7 | -8.8 | **-12.6** |
| Nb03 | -10.3 | -10.1 | **-11.6** | -11.9 | **-15.9** | -5.8 | -8.9 | -3.8 | -7.0 | -8.9 |
| Nb04 | -4.0 | -8.7 | -10.9 | -7.3 | -18.0 | -10.2 | -9.2 | -8.6 | **-12.4** | -11.0 |
| Nb05 | -11.6 | -8.8 | -10.8 | -11.2 | -13.5 | **-13.9** | -7.1 | -8.5 | -7.2 | -3.5 |
| S43 | -6.6 | -7.4 | -9.2 | -11.1 | -9.5 | -5.2 | -6.0 | -5.1 | -10.9 | -9.8 |
| bn03 | -12.0 | -7.3 | -8.5 | -10.8 | -8.8 | -9.3 | -4.7 | -8.1 | -4.0 | -3.5 |
| R14 | -9.7 | -6.8 | -7.3 | -10.3 | -7.7 | -3.5 | -4.0 | -2.1 | -3.5 | -10.9 |
| 3-2A2-4 | -4.2 | **-11.3** | -6.7 | -6.3 | -9.5 | -7.7 | -6.9 | -4.7 | -4.5 | 0.9 |

**TABLE VI.**
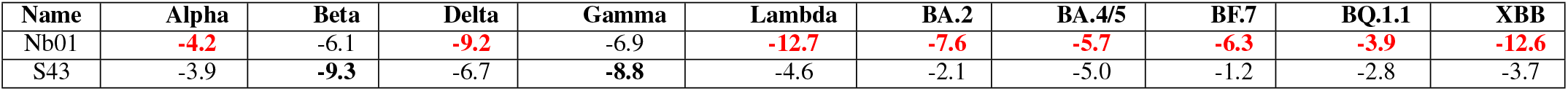
Affinity of Nb01 and s43 to compete for binding to the same SARS-CoV-2 variant (kcal/mol)

The reason to explain the strong binding affinity of Nb01 shows as follow. *Hydrogen bonds formed in the Nb01 CDR3 region is much stable in competition experiments*. Figures 6A show the binding details of Nb01, and S43. Clearly, the binding epitopes of Nb01 and S43 are very close to each other, with the binding pocket located between amino acids 100 and 200 of the variant 3CL^pro^. The residues 80 to 90 of Nb01, which is the CDR3 region of Nb01, are crucial for the affinity to the variant, primarily through the formation of hydrogen bonds in this region. Extended Data Table I-XX show the interaction forces of Nb01 and S43 when bound to SARS-CoV-2 variants, respectively. Extended Data Table XXI-XL shows the interaction forces generated when Nb01 and S43 compete to bind the same variant. Extended Data Figure 1 shows the residues necessary for variant binding when competing with each other. The results show that Nb01 have more stronger hydrogen bonds than S43 in CDR3 area.

**Fig. 6.**
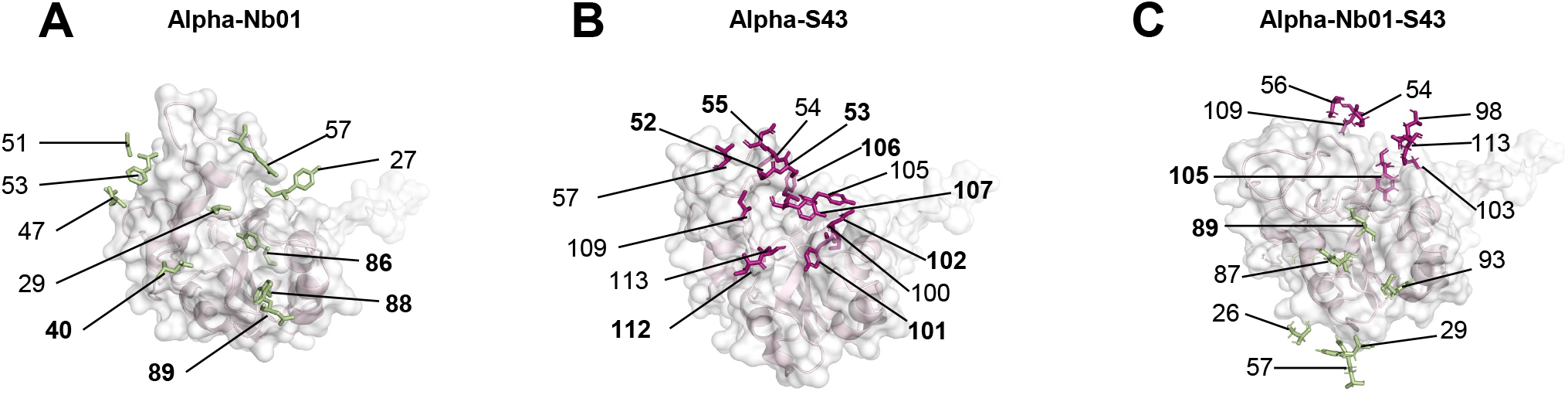
Alpha complex interaction details. (A) Interaction details of Nb01 and Alpha complex; (B) Interaction details of S43 and Alpha complex; (C) Interaction details of Nb01 and S43 upon simultaneous binding to Alpha. Bolded parts are residues that generate interaction forces. Residues that are crucial for interaction are numbered. “ 1” denotes the N-terminal residue of the nanobody.

#### 3) Complexes of Nb01 with SARS-CoV-2 variants are more stable

*Nb01 complex has the lower potential energy than S43 complex*. The potential energy changes of Nb01 and S43 is calculated by molecular dynamics simulation when they bind to the SARS-CoV-2 variants. Table VII shows that Nb01 has the lowest potential energy (the average potential energy is 746) with the various SARS-CoV-2 variants, which indicated that Nb01 is the most stable upon binding (Extended Data Figure 2).

**TABLE VII.** Potential energy changes in complexes of Nb01 and S43 with variants.

| Name | Alpha | Beta | Delta | Gamma | Lambda | BA.2 | BA.4/5 | BF.7 | BQ.1.1 | XBB |
| --- | --- | --- | --- | --- | --- | --- | --- | --- | --- | --- |
| Nb01 | **750** | **800** | **560** | **700** | **900** | **700** | **750** | **800** | **800** | **700** |
| S43 | 850 | 850 | 650 | 800 | 1000 | 800 | 800 | 900 | 900 | 800 |

## IV. Conclusion

We selected 3CL^pro^, the major protease of COVID-19, as the target protein, generated Nb01 with high activity towards 3CL^pro^, and verified the properties of Nb01 by molecular docking and molecular dynamics simulations. The results showed that Nb01 was excellent in terms of binding affinity (approximately twice the baseline) and other physicochemical parameters (GRAVY = −0.17, Toxins *<* 0.6, Instability Index = 41.52 and Aliphatic Index = 72.22). In addition, the generated nanobody Nb01 is active against most of the current SARS-CoV-2 variants. Nb01 is active against most of the current SARS-CoV-2 variants, it possesses the lowest binding energies for Alpha, BA.4/5, and BF.7, and its affinity for other variants is comparable to other nanobodies. The higher stability of Nb01 compared to the well-known nanobody S43 suggests that Nb01 is an excellent lead compound for the SARS-CoV-2 variant. In future, we will perform in vivo and vitro experiments on Nb01.

## Acknowledgment

This research work was financially supported by Shenzhen Second People’s Hospital COVID-19 Emergency Clinical Research Project(2023xgyj3357009) and Shenzhen Science and Technology Program(20231127194506001), Shenzhen Science and Technology Innovation Committee Funds (JSGG20220919091404008), Shenzhen Technology University Innovation and Entrepreneurship Project (S202414655005) and the National Natural Science Foundation of China (81804154).

